# On the robustness of scRNA-seq foundation models for plant perturbation response prediction under cross-experiment shift

**DOI:** 10.64898/2026.08.21.746324

**Authors:** M. Fernandez Burda, R. Bonazzola, A.A. Valli, G. Castrillo, G. Stegmayer, E. Ferrante, D.H. Milone

## Abstract

Foundation models for single-cell transcriptomics promise to learn generalizable representations of cellular states. However, recent evidence suggests they often fail to outperform simple machine learning baselines. Furthermore, their ability to generalize across unseen experimental conditions remains poorly understood, particularly in plants, where rigorous evaluation beyond cell type annotation and batch integration is lacking. To address this, we introduce an *Arabidopsis thaliana* foundation model, scAraFM, and benchmark it across several perturbation conditions under three increasingly challenging protocols: random splits from a single experiment, replicate-based splits, and cross-experiment transfer learning. We found that random splits overestimate performance by up to 30 points relative to cross-experiment evaluations. Across representation strategies, preserving gene identity consistently outperforms the standard pooled embeddings. Moreover, simple baselines using raw reads remain competitive in single-experiment settings, challenging current claims of universal advantage of foundation models. In contrast, under cross-experiment transfer, pretrained representations show added value, particularly with few labelled samples, suggesting that the benefits of foundation models emerge precisely in the regimes that matter for practical deployment. Overall, our results demonstrate that conclusions about foundation models depend critically on the evaluation design, and that preserving per-gene structure aids generalization in downstream tasks, supporting robust predictions across unseen experimental contexts.

## Introduction

The advent of single-cell omics has changed our understanding of biological systems, offering a detailed view of cells and their functional dynamics. This includes insights into cell types, states and their changes during development, disease and therapeutic response. The vast volume and heterogeneity of the data generated presents both opportunities and challenges. Current analytical methods fall short in capturing variation across diverse large-scale single-cell datasets, for example when experimental conditions differ, motivating the development of novel computational strategies. In parallel, the artificial intelligence community was revolutionized by the introduction of Transformer architectures^1^. Although initially designed for natural language processing, they are now being pretrained on large amounts of domain-specific data (such as biological sequences), and used as feature extractor of machine learning (ML)-based methods. These pretrained stages are known as foundation models (FMs), and can be effectively adapted to a wide range of bioinformatics tasks with additional fine-tuning^2–4^.

Foundation models have quickly become the standard approach for representing single-cell transcriptomes. Examples of single-cell FM (scFM) include scBERT (2022), which demonstrated strong performance on cell-type annotation and subsequent work on transcriptomic language modelling^5^; Geneformer (2023), pretrained on single-cell transcriptomes to enable transfer learning for context-specific predictions in network biology and related tasks^6^; scGPT (2024), a generative pretrained transformer trained at atlas scale and used for downstream tasks such as cell-type annotation, data integration, and multi-omic modeling^7^; and scFoundation (2024), a large-scale pretrained model designed to capture gene to gene context relationships across diverse cell types and states^8^. However, FMs available for transcriptomics are mostly centred on human and mouse species, mainly because of the large amount of available training data. For instance, Geneformer and scGPT were trained on tens of millions of primarily mammalian single-cell profiles, and scFoundation was pretrained on more than 50 million human single-cell transcriptomes. By contrast, plant single-cell resources are expanding but remain much smaller and more heterogeneous, with an order-of-magnitude less data relative to mammalian pretraining corpora^9,10^. Two recent efforts have introduced dedicated scFMs for *A. thaliana*: scPlantLLM^10^ and scPlantFormer^11^, both pretrained on approximately one million *A. thaliana* cells. However, their evaluation has largely been limited to cell-type annotation and batch integration, leaving unanswered the question of whether plant scFMs can generalise to more biologically challenging tasks such as perturbation response prediction or stress classification, which are the focus of the present study. Collectively, these models are used as general-purpose backbones: pretrained once, and then adapted with minimal supervision to a broad variety of single-cell specific tasks.

Several benchmarking efforts have attempted to quantify in which scenarios scFMs actually add value over classical pipelines. For instance, the evaluation of scBERT for cell-type annotation was replicated^12^, Geneformer and scGPT were assessed in zero-shot settings for cell-type structure and batch integration^13^, and complementary benchmarks have focused on perturbation-response prediction where testing data distribution is different from the training data (i.e. distribution shift)^14,15^. At a broader level, it was argued that simple ML models can adapt more efficiently to specific datasets, and that no single scFM consistently dominates across multiple tasks, implying that gains are task- and dataset-dependent^16^. Across these works, a recurrent conclusion has emerged: pretrained scFMs often fail to outperform simpler baselines. Multiple foundation models for perturbation prediction were also bench-marked^17^, finding that none of them outperformed simple linear or additive baselines, while it was also reported that scFM embeddings do not yield consistent gains, especially under distribution shift^18^. Together, these studies motivate doubts about the performance gains of representations learned by scFM, when compared to more plain alternatives, and highlight the need for protocols that explicitly test robustness and generalization under more complex tasks.

In this work, we establish a standardized evaluation framework with increasing levels of challenge in the prediction of plant perturbation responses, to assess when FMs offer genuine advantages with respect to classical ML approaches. Concretely, our evaluation design escalates from (i) within-study random splits, training and testing on the same dataset at random; to (ii) replicate-aware splits, holding out biological or technical replicates; and finally to (iii) cross-Experiment evaluation, where development and testing occur on independent studies targeting the same condition. These increasingly complex evaluation scenarios force models to confront broader generalization regimes. To address the reported major issues and extend the analysis, it is required to pretrain the scFM for each evaluation scenario excluding the data that will be used in the subsequent downstream validation stage. Thus, we developed a plant transcriptomic FM pretrained on *A. thaliana* single-cell data, named scAraFM.

Results show that evaluation settings that approximate real testing scenarios have a concrete difficulty, which is overestimated in within-study random splits. At the FM architectural level, we have also seen that representations that preserve gene identity are more robust than just a condensed single vector for each cell. Moreover, representations of scFM do not reliably outperform strong classical baselines trained directly on read counts; that is, raw-read baselines exceed in performance, except in low data regimes for cross-experiment evaluation, where FMs provide added value. Finally, we establish an explicit stress-testing framework for transcriptomic embeddings in plants, emphasizing that claims of generalization should rely on replicate-aware or cross-experiment evaluation rather than random data splits. We envision that this validation framework will serve as a standard for future transcriptomic models, providing clear metrics for when and where FMs deliver value.

## Results

We developed scAraFM, a transformer-based single-cell FM pretrained on *A. thaliana* scRNA-seq data. We assembled publicly available datasets and curated tissue-specific pretraining corpora comprising approximately 17 M cells drawn from 26 independent studies (Supplementary Table 1). Counts were library-size normalized and log-transformed. Highly variable genes (HVGs) were identified per dataset and a consensus set of genes was constructed via frequency-based ranking across datasets (Figure 1a; see full details in Methods). Architecturally, scAraFM follows a scBERT-style^5^ encoder in which genes are treated as tokens and cells as sequences. Each gene is mapped to a learned embedding vector, and normalized expression values modulate the corresponding gene embeddings before being passed through the transformer encoder. The model is pretrained by predicting masked gene tokens from the remaining observed genes, to learn context-aware gene representations that capture co-expression patterns and high-order dependencies (Figure 1a, right). Throughout downstream benchmarking, scAraFM was used as a frozen feature extractor: embeddings were computed once per cell and subsequently processed by lightweight supervised classification heads, allowing the evaluation to focus directly on how different representation choices perform under increasing distribution shift. Importantly, for each evaluation scenario, scAraFM was pretrained while explicitly excluding the data later used in the corresponding downstream evaluation stage.

**Figure 1:**
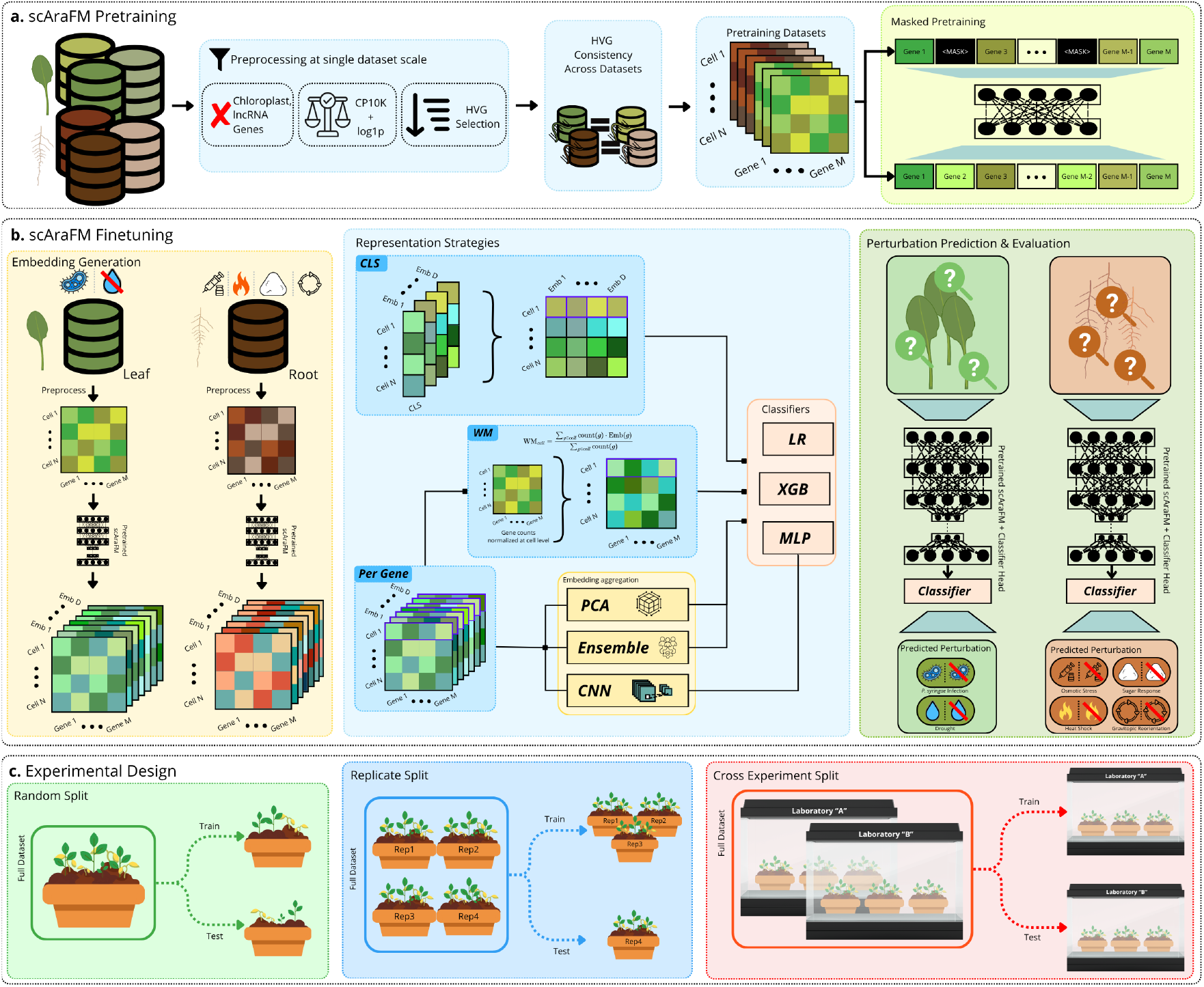
scAraFM training and evaluation overview. (a) scAraFM pretraining. Raw *A. thaliana* scRNA-seq datasets are processed independently at the single-study level: gene identifiers are harmonized, organellar genes and lncRNA are removed, counts are normalized (CP10K) and log-transformed (log1p), and highly variable genes (HVGs) are selected. HVG sets are then reconciled across datasets. The foundational model (in the right) is trained with masking self-supervision, predicting masked gene tokens from the remaining observed genes to learn context-aware gene representations. **(b) scAraFM finetuning**. The pretrained scAraFM encoder produces per-gene embeddings. For each labeled dataset, the same preprocessing is applied ensuring that the same gene subset is utilized in the pretraining phase. Multiple representation strategies are evaluated (in the center), using CLS token pooling (top), an average of gene embeddings weighted by expression counts (WM, middle), or the full per gene embedding (Per Gene, bottom). We highlight in purple a single cell sample to better visualize dimensionality of each representation. These are fed to downstream predictors: logistic regression, gradient-boosted trees (XGB) and CNN/MLP, with exception of the per-gene strategy that is first processed to reduce gene dimensionality before reaching the classifier heads (PCA and ensemble). The joint encoder-classifier is trained and evaluated on condition prediction tasks (right): *P. syringae* infection and drought for leaves, and osmotic stress, sugar response, heat shock and gravitropic reorientation for roots. **(c) Experimental Design**. We evaluated three train-test protocols with increasing generalization difficulty: (i) random split (left), where train/test splits are drawn at random from the same dataset; (ii) replicate-aware split (center), where an entire biological replicate is held out for testing (and the remaining replicates are used for training); and (iii) cross-experiment split (right), where the model is trained on one dataset and evaluated on an independent dataset generated in a separate experiment.

### Increasing difficulty grades emerge across generalization splits

A central question in applying FMs to downstream tasks is how to transform high-dimensional per-gene embeddings into a representation suitable for classification. The strategies that compress an entire cell into a single vector are convenient, but they may discard inter-gene patterns of expression that prove critical for downstream tasks. To systematically explore this trade-off, we evaluated five strategies derived from scAraFM embeddings alongside a baseline with raw normalized counts, each paired with multiple classifier heads (Figure 1b, center). At the cell level, two compression strategies were considered, also known as pooling schemes in deep learning: classification token (CLS), which uses the final-layer embedding; and gene-expression weighted mean (WM) of the embeddings. At the gene level, the full gene-by-embedding matrix was retained and processed through three alternatives: PCA over the embedding dimensions; ensemble over de embeddings dimensions (Ensemble); and a 1D convolutional layer (CNN) that aggregates the embedding dimensions. Then, extracted features feed a logistic regressor (LR), an extreme gradient boosting classifier (XGB), and a multi-layer perceptron (MLP). To benchmark these representations, datasets were curated covering 7 perturbation conditions (Figure 1b right and Supplementary Table 2). Classes were defined as binary, that is, having/not having the perturbed condition, and the performance was assessed by the area under the receiver operator curve (AUROC) (see Methods and Appendix B). Finally, to test whether the evaluation protocol itself influences the conclusions drawn about representation quality, three train-test split regimes were designed with increasing generalization difficulty. In the within-study random split (Figure 1c, left), training and test sets are drawn at random from the same dataset; this is the standard protocol often used in literature^19–21^. In the replicate-aware split (Figure 1c, center), an entire biological or technical replicate is held-out for testing, so that train and test share the same experimental context but contain no overlapping replicate identifiers. In the cross-experiment split (Figure 1c, right), the model is trained on one dataset and evaluated on an entirely independent dataset targeting the same perturbation condition, thus testing transfer across independent studies.

Applying these protocols to the perturbation classification task (Figure 2a) reveals a clear and consistent difficulty gradient. Performance for all methods is highest under the random split, drops when a replicate is held out, and declines most sharply under the cross-experiment split, where evaluation occurs on an entirely independent study. This ordered degradation (random ¿ replicate ¿ cross-experiment) supports the hypothesis that increasing shifts in experimental sourcing introduce distributional changes that challenges the generalization capability of the models. Quantitatively, random splits overestimate performance by 20–30 AUROC points relative to replicate-held-out evaluation and by an even wider margin relative to cross-experiment transfer, underscoring the need for study-aware validation of FMs in transcriptomic data. The cross-experiment setting, although constrained by the limited availability of public datasets with compatible perturbations, provides the strictest and most reliable assessment of real-world generalization in perturbation prediction. These results indicate that conclusions drawn from within-study random splits can be overly optimistic, and that claims about representation quality should be validated against protocols that better approximate realistic conditions.

**Figure 2:**
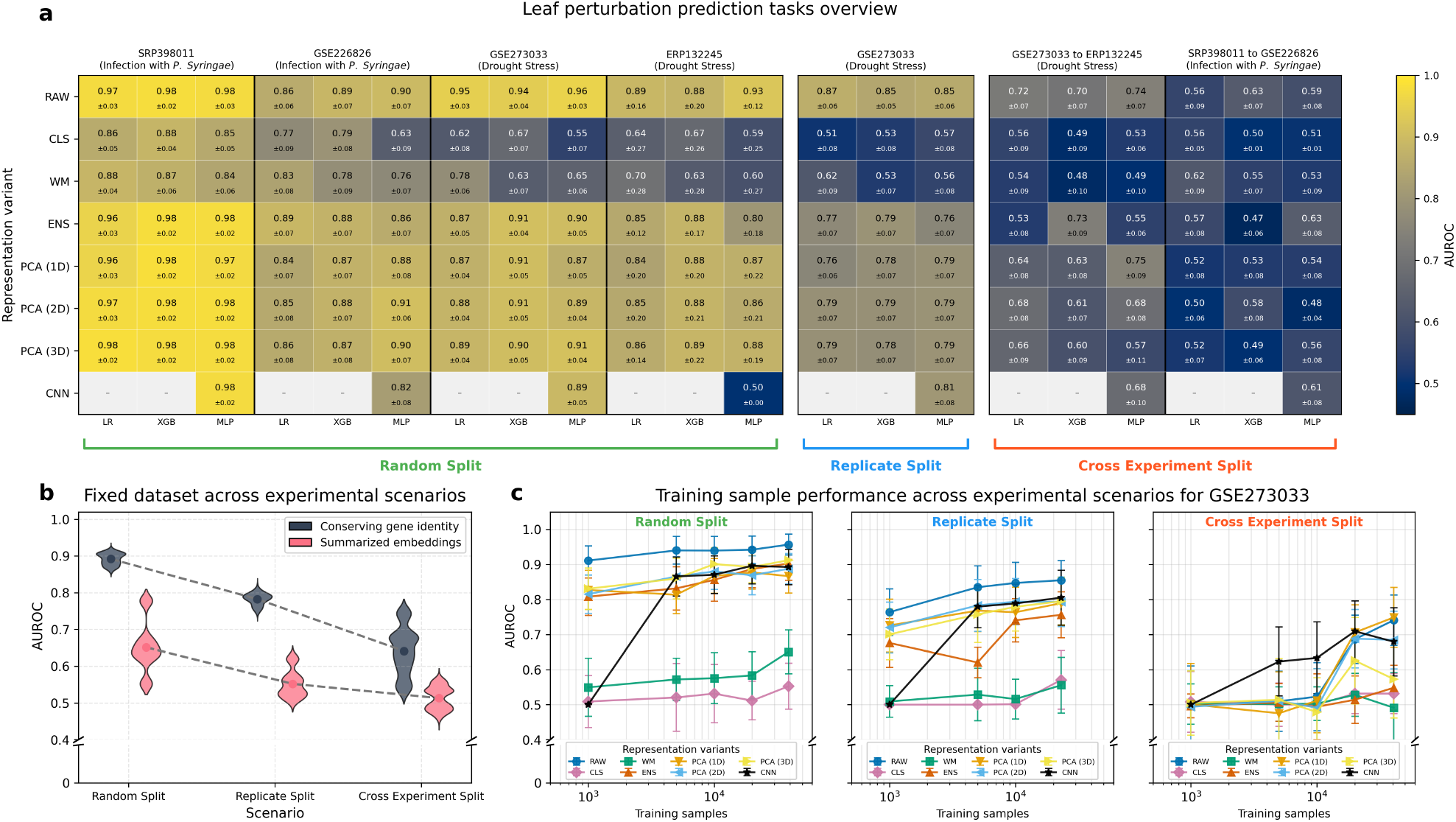
Perturbation prediction performance on varying scenarios for *A. thaliana* leaves datasets. a) Each heatmap cell depicts AUROC for all classifiers (LR, XGBoost, and MLP) along the representation variants (RAW, CLS, WM, ENS, PCA and CNN). On the columns, the classification tasks are shown in rising difficulty, from left to right: random splits, replicate splits and cross-experiment splits. Each perturbation condition is indicated below the dataset ID. The standard deviation (± SD) of the metric is obtained from 100 bootstrap resamples of predictions in the test set. b) AUROC distributions on different evaluation scenarios, from random splits (left) to cross-experiment split (right), when using gene-identity conserved embeddings versus pooled summarized embeddings. c) Performance of the MLP classifier for every representation variant as the number of training samples increases. These increasingly challenging experimental scenarios are shown for the GSE273033 dataset (drought stress)

### Preserving gene identity is crucial for robust classification in cross-experiment transfer

An important aspect of the evaluation is the assessment of the impact of summarizing gene information versus preserving gene identity. To this end, performance of representation variants can be considered divided into schemes condensed at cell level (WM and CLS) versus approaches that retain the gene identity (ENS, PCA, and CNN). Figure 2b shows that, across all three evaluation regimes in leaf tissue, a consistent pattern emerges: representation strategies that preserve gene identity at the input of the classifiers outperform those that compress gene information into a single summarized vector. In particular, the schemes condensed at cell level systematically underperform, whereas approaches that retain gene identity achieve consistently stronger results. This pattern holds regardless of the down-stream classifier employed (LR, XGBoost, or MLP), suggesting that it reflects a genuine property of the representation rather than an artefact of classifier choice. Maintaining a gene-level view of the cell likely enables classifiers to exploit gene-gene dependencies and interaction structure in the latent space, which are blurred or lost under aggressive summarization. This is primarily because, unlike words in a sentence, in scRNA-seq data all genes are always present at the input, and in the same position. Therefore, what is relevant for downstream tasks is not the position itself of the gene, but its expression level relative to other genes in the same cell.

The results in Figure 2a (right) showed that cross-dataset transfer was the hardest evaluation scenario and that the limiting factor is the distribution shift: models trained on one experiment struggle when the data-generating process changes. At maximum available training data, gene identity preserving strategies achieve the best or tied-best performance across classifiers on cross-experiment splits: ENS & CNN on *P. syringae* infection, and PCA (1D) on drought stress (detailed on the cross experiment snippet of Figure 2a). Moreover, this shift can have more impact when data size changes. Within this harder regime, scAraFM representations provide robust transfer learning in the low-data regime (Figure 2c, right), with the CNN+MLP classifier standing out as a particularly effective combination, outperforming from low data to full data regimes all other input types and models. More broadly, classifiers based on pretrained embeddings tend to saturate close to the performance achieved by direct transcriptomic baselines. In summary, scAraFM results show that representation learning can improve robustness under distribution shift, especially on the cross-experiment scenario and in low-data regimes, where the CNN+MLP finetuning approach gains a differential performance when compared to all other methods in the early 5,000 and 10,000 training samples (Figure 2c, Supplementary Figure 1 and 2).

An important point is that under random and replicate splits scenarios, simple baselines using raw transcriptomic reads remain competitive to learned embeddings. LR or XGB trees with raw input data frequently match the performance of models with input embeddings at a fraction of the computational cost. We interpret this not as a refutation of representation learning, but as evidence that the added value of FM representations materializes most clearly under distribution shift, precisely the regime that matters for practical deployment. Pretrained embeddings appear to provide a more efficient feature space than raw counts in cross experiment and low data regimes, reducing the number of labelled cells required to reach a given performance level, which is especially relevant in plant scRNA-seq where many conditions of interest are represented by small studies and sparse annotation.

## Discussion

This work addresses two important and interrelated aspects of transcriptomic representation learning in plants, using *A. thaliana* single-cell RNA-seq as a concrete testbed. First, evaluation protocols that better approximate real-world use cases, by holding out replicates and entire experiments, induce a consistent difficulty gradient for assessing models performance. This pattern supports the hypothesis that random splits within a dataset can overestimate performance by allowing specific effects of a study (batch, protocol, and label-correlated confounders) to persist across train and test. Second, representations that preserve gene identity, that is, each gene with its own separated features, are systematically more robust than representations that condense a cell into a single summarized vector. This pattern is reproduced in root tissue across five additional perturbation prediction tasks (osmotic stress, heat shock, sucrose response, estradiol induction and gravitropic reorientation) where gene-identity-preserving variants again exceed raw-count baselines while pooled summaries underperform (Supplementary Figure 3). The consistency across two tissues and seven distinct perturbations supports interpreting this as a property of the representation itself. Our results are consistent with recent evidence in the broader scFM literature that features at gene resolution can capture co-regulatory modules and expression signatures more faithfully than ad-hoc cell summaries^16^. Finally, beyond reporting model comparisons, we provide an explicit framework of increasing difficulty for transcriptomic embeddings, emphasizing that claims about “generalization” should be tied to replicate-aware and cross-experiment evaluation rather than random splits inside the datasets.

A practical implication of the above is that the choice of representation is not only a question of accuracy, but also of computational budget and iteration speed. Training and maintaining transcriptomic FMs can be computationally intensive: large-scale pretraining requires substantial GPU resources, careful engineering, and repeated experimentation over architecture, tokenization and normalization. By contrast, classical models trained directly on transcriptomic profiles are comparatively easier to develop, often feasible on modest hardware, and enable rapid baseline iteration. In our experiments, these simple models frequently saturate performance quickly as training size increases, explaining why they remain competitive under within-study evaluations. However, when exposed to cross-experiment settings, they exhibit the same limited generalization capacity as more complex models. Ultimately, strong raw baselines are an essential reference point that prevents over-attributing gains to representation learning when similar performance is attainable with far lower computational cost.

The evaluation setup proposed here also makes clear where methodological progress is still needed. The cross-experiment setting is the strictest test of whether a representation captures transferable biological signals rather than specific artifacts of an experiment. In that regime, simple classifiers based on raw reads can degrade to near-chance performance, suggesting that the limiting factor is the distribution shift: models trained on one experiment lower performance when the protocol of data generation changes. While performance gains are promising, the sensitivity under cross-experiment transfer highlights a valuable opportunity for developing methods that explicitly target robustness to protocol and batch variation, improve alignment across studies, and learn invariances that persist across independently generated datasets. Our findings therefore align with the broader message that within-dataset performances are insufficient evidence of meaningful generalization, echoing prior observations that simple baselines may rival large transcriptomic models under random splits^12^. Therefore, we interpret these results as a call to measure performances in regimes closer to real world scenarios: transfer learning to unseen studies. Crucially, while releasing FMs pretrained on all available data is a standard practice, rigorously evaluating generalization capacity requires pretraining datasets to strictly exclude all data used for downstream validation tasks, particularly in cross-experiment benchmarks.

At the same time, a recurring pattern is that pretrained representations can be beneficial in low data regimes, where embeddings may provide a more efficient feature space than raw counts, reducing the number of labeled cells required to reach a given performance level. This potential advantage is especially relevant in plant scRNA-seq, where many conditions of interest are represented by small studies and sparse annotation. These findings are grounded in the framework of the present study: the available single-cell data and perturbation prediction tasks for *A. thaliana*, which define the scope of our observations. Plant scRNA-seq resources remain smaller and more heterogeneous than human atlases, limiting both pretraining scale and the diversity of protocols, genotypes, and perturbations available for robust transfer tests. Progress will likely require (i) larger, better-curated corporas (ideally multi-species), benchmark designs that privilege replicate-held-out and cross-experiment evaluation as defaults, and (iii) representation strategies that preserve gene identity while being explicitly robust to experimental variation. Taken together, our results motivate continued development of transcriptomic FMs, but with an empirical standard that rewards cross-study transfer: if the goal is reusable representations, the evaluation should mirror this utilization.

## Conclusions

This study provided a rigorous evaluation design for transcriptomic FM representations in the single-cell domain, explicitly prioritizing evaluation regimes that better reflect real-world use. By separating random splits within each dataset, replicate held out, and cross-experiment scenarios, a consistent difficulty gradient was revealed, demonstrating that conclusions drawn from within-dataset random splits can be overly optimistic due to specific conditions of the study. Specifically, our results highlight two key insights regarding representation choice: first, representations that preserve gene identity are more robust than cell summaries; second, pretrained representations provide a significant advantage when task-specific training data is scarce, underscoring the role of foundation models as reliable feature extractors, preferred over simpler approaches. While our claims were demonstrated using *A. thaliana* datasets, the remaining performance gap in cross experiment transfer is an open call for methods that explicitly target robustness to protocol and batch variation. More broadly, we argue that cross-study evaluation should become the default standard for assessing transcriptomic embeddings, and we hope this study helps steer future work toward specific corpora, more biologically structured representations, and evaluation protocols that reward transferable signals.

## Methods

### Data preparation

A compendium of publicly available *A. thaliana* transcriptomic datasets was assembled (sc-PlantDB^22^, NCBI GEO^23^ and SRA^24^), comprising approximately ~ 834 k leaf cells and ~ 878 k root cells, and prioritizing single-cell RNA-seq studies with well-annotated experimental metadata. To support downstream benchmarking, a subset of datasets covering diverse perturbations was also curated (e.g., drought, pathogen infection, osmotic stress, heat shock, sugar response and gravitropic reorientation), stratified by tissue (leaf vs. root) and organized into the evaluation protocols described in the experimental design (Supplementary Table 1).

All scRNA-seq datasets were processed with a standardized Scanpy-based pipeline to ensure a consistent feature space across studies and to make cross-dataset comparisons well-posed. Briefly, gene identifiers were harmonized to canonical *A. thaliana* AGI locus identifiers, collapsing duplicated entries and filtering non-comparable identifiers (chloroplasts, lncRNAs). Raw counts were then normalized per cell to a fixed library size followed by log-transformation. Within each dataset, HVGs were identified using Scanpy^25^ (Seurat flavor), and then aggregated into a cross-dataset consensus set. The individual steps of this preprocessing pipeline are detailed below.

1. **Gene identifier harmonization and filtering**. Gene identifiers were first harmonized to the *A. thaliana* locus nomenclature. Duplicate gene entries in the expression matrix were collapsed by summing counts across duplicated rows, yielding a one-gene to one-feature representation. Features were filtered to retain only canonical *A. thaliana* locus identifiers. To avoid expanding the feature space with dataset-specific or non-comparable annotations, organellar loci and non-target identifiers were excluded from the feature set. This exclusion applied to chloroplast-prefixed loci such as ATC, and long non-coding RNA identifiers such as ATHLNC.
2. **Library-size normalization and log transformation**. To correct for sequencing-depth differences across cells, raw counts were normalized on a per-cell basis to a fixed library size (counts per 10, 000) using Scanpy’s total-count normalization, followed by log-transformation with log1p(*x*) = ln(*x* + 1).
3. **Highly variable gene discovery**. Within each dataset, HVGs were identified using Scanpy’s dispersion-based procedure with the Seurat flavor, which bins genes by mean expression and ranks them by normalized dispersion within bins to prioritize biologically informative variation while controlling for mean-variance dependencies.
4. **Consensus HVG construction**. To enable cross-dataset analyses in a shared feature space, HVGs were aggregated into a consensus list using a frequency-based criterion: genes were ranked by the number of datasets in which they were selected as HVGs, and the top genes were retained. This approach emphasizes genes that are reproducibly variable across studies rather than dataset-specific artifacts.

### scAraFM and representation strategies

scAraFM was used as a frozen feature extractor for *A. thaliana* single-cell transcriptomes, following a scBERT-style transformer encoder in which genes are treated as tokens and cells as sequences. To ensure that both pretraining and downstream evaluation operate in a shared and comparable feature space (Figure 1.a), all datasets are processed with a consistent pipeline: gene identifiers are harmonized to canonical *A. thaliana* locus names, duplicated loci are collapsed by summing counts, and non-comparable identifiers (e.g., organellar loci and dataset-specific annotations) are removed. Counts are library-size normalized (counts per 10,000 cells) and log-transformed, and a consensus HVG set was built by ranking genes by how frequently those are selected as HVGs across datasets and retaining the top *G* = 4, 000 genes. This yields a fixed gene vocabulary and a consistent input dimensionality across studies. Architecturally, scAraFM maps each gene *g* to a learned embedding vector in ℝ^*D*^, with *D* = 200. For a cell *c*, normalized expression values *x*_*c,g*,_ are used to modulate the corresponding gene embeddings before being passed through the transformer encoder, producing contextualized, gene-specific representations that capture co-expression patterns. We pretrain tissue-specific instances of scAraFM on curated *A. thaliana* corpora collected from 26 public datasets (Supplementary Table 2). Throughout our downstream perturbation-prediction benchmarks, we treat scAraFM primarily as a representation model: embeddings are extracted once per cell and then consumed by lightweight supervised heads, allowing evaluation to focus on how representation choices behave under increasing distribution shift.

Let *h*_*c,g*_ ∈ ℝ^*D*^ denote the final-layer embedding produced by scAraFM for gene *g* in cell *c*. We compare representation strategies that differ in how they compress cell information (Figure 1.b).

### Cell-level representations

1. **[CLS] pooling (CLS)**. We use the final-layer [CLS] embedding as a single-vector representation of the cell *h*_*c,CLS*_. This is a compact option (of dimension *D*) that mirrors standard transformer aggregation.
2. **Expression-weighted mean (WM)**. We compute an expression-weighted average of the gene embeddings; *WN* (*c*) = ∑_*g*_ *x*_*i,g*_*h*_*c,g*_*/* ∑_*g*_ *x*_*c,g*_, where *x*_*i,g*_*′ /*∑ _*g*_ *x*_*c,g*_ is the normalized expression of gene *g*^*′*^ in cell *c*. WM keeps an explicit coupling between transcript abundance and the embedding while still producing a single cell vector of size *D*.

#### Gene-level representations

3. Instead of collapsing gene information into one vector, we retain the gene-by-embedding matrix *H*_*c*_ = [*h*_*c*,1_*h*_*c*,2_ · · · *h*_*c,G*_] ∈ ℝ^*D×G*^ and adapt it for downstream classifiers using three alternatives.
  3.1 **PCA over embedding dimensions (PCA)**. We apply PCA to reduce the embedding dimension from *d* to a small number of components *k* (here *k* ∈ {1, 2, 3}), fitting PCA on the training set and applying the same projection to development and test sets. After flattening out, this produces a compact representation in ℝ^*G×k*^ that retains gene identity but removes redundancy across embedding channels.
  3.2 **Embedding-dimension ensemble (ENS)**. For each embedding coordinate *d* ∈ {1, …, *D*}, we form a gene-level feature vector by taking the *d*−th component across all genes in *H*_*c*_. We train one classifier per coordinate and aggregate predictions across coordinates by majority vote. This preserves gene identity while decomposing the full tensor into simpler views, reducing the effective dimensionality seen by any single classifier to a vector of size *G*.
  3.3 **Convolutional head over per-gene embeddings (CNN)**.Mirroring finetuning in scBERT original implementation^5^, we train a lightweight convolutional classifier that consumes *H*_*c*_ directly. A 1D convolutional module compresses and mixes information across the embedding dimension, followed by a shallow MLP to produce predictions. This approach allows task-adaptive aggregation while keeping gene identity explicit through most of the computation.

For all representations except the CNN head, which defines its own classifier, we pair the extracted features with standard supervised heads. Therefore, logistic regression, MLP, and XGBoost heads were trained on the perturbation tasks.

### Experimental Design

Models were benchmarked in three scenarios on the perturbation classification task (Figure 1.c). These experimental conditions underlie an increasing difficulty spectrum as they incrementally test broader generalizability of the benchmarked methods.

#### Common pipeline

For each dataset, the following steps were applied:

i. Preprocessing. From the expression matrices, the preprocessing ensures a consistent feature space and condition labeling across experiments.
ii. Class balancing. Before defining splits, data was balanced by class by downsampling each class to the size of the least represented class. This yields a class-balanced pool used for all subsequent split procedures.
iii. Experimental split definition. Train/test partitions are built according to the split protocol (Random, Replicate, or Cross-Experiment; detailed below). When applicable, the test set is capped (*N* ≤ 2, 000) to avoid excessively large evaluations and to keep results comparable across datasets.
iv. Training sample capping. From the training partition, we run learning curves by capping the total samples (whenever possible) to values of 1,000, 5,000, 10,000 and 20,000 maintaining the same fixed test set. Additionally, we perform a final training setup with all available training data after holding out the test data. All the training and test sets created in these setups are class-balanced.
v. Model training and evaluation. For each representation variant and classifier, we train on the limited training set and report metrics on the fixed test set for that experimental configuration.

#### Random split

A class-balanced random split was performed, with a test size of 20% of the balanced pool. To prevent overly large test sets, we apply the following rule. Let *N* be the total number of samples after class balancing.

- Test set: Define the target test size as 0.2*N*. If 0.2*N >* 2, 000, we limit the test set to 2, 000 samples, using stratified sampling to preserve class balance. Otherwise, we use the full 0.2*N* balanced test set.
- Training set: all remaining balanced samples, then limited by training count (or ALL).

#### Replicate split

We split by biological/technical replicate to prevent overlap between replicates across train and test.

- Test set: the first replicate (according to the replicate ordering defined in the dataset metadata). If the test replicate contains more than 2, 000 samples, we limit it to 2, 000 using stratified sampling by class.
- Training set: all remaining replicates (from the balanced pool), subsequently subject to the training-count limiting described above.

#### Cross-experiment split

We evaluate cross-dataset transfer by training on one dataset and testing on another dataset that shares the same label space (i.e. downstream task):

- Test set: an entire target dataset, after applying the same preprocessing and class balancing procedure. If the target dataset has more than 2, 000 samples, we limit the test set to 2, 000 using stratified sampling by class.
- Training set: the source dataset (balanced), then limited by training count (or ALL).

## Author contributions statement

D.H.M. and E.F. conceived the study and designed the methodology. D.H.M. led the project. R.B. and M.F. curated the data, wrote the code and performed experiments. M.F. prepared figures. D.H.M., E.F. and G.S. supervised the work and analyzed the results. A.A.V and G.C. provided biological expertise and interpretation. G.C., A.A.V., G.S., D.H.M., and E.F. acquired funding through joint grant application. M.F., D.H.M. and G.S. co-wrote the original draft of the manuscript. All authors reviewed and approved the manuscript.

